# Pheromone representation in the ant antennal lobe changes with age

**DOI:** 10.1101/2024.02.13.580193

**Authors:** Taylor Hart, Lindsey E. Lopes, Dominic D. Frank, Daniel J.C. Kronauer

**Affiliations:** Laboratory of Social Evolution and Behavior, The Rockefeller University, 1230 York Avenue, New York, NY 10065, USA; Howard Hughes Medical Institute, New York, NY 10065, USA

**Keywords:** age polyethism, calcium imaging, clonal raider ant, communication, division of labor, odor coding, olfaction, *Ooceraea biroi*

## Abstract

While the neural basis of age-related decline has been extensively studied (*1–3*), less is known about changes in neural function during the pre-senescent stages of adulthood. Adult neural plasticity is likely a key factor in social insect age polyethism, where individuals perform different tasks as they age and divide labor in an age-dependent manner (*4–9*). Primarily, workers transition from nursing to foraging tasks (*5*, *10*), become more aggressive, and more readily display alarm behavior (*11–16*) as they get older. While it is unknown how these behavioral dynamics are neurally regulated, they could partially be generated by altered salience of behaviorally relevant stimuli (*4*, *6*, *7*). Here, we investigated how odor coding in the antennal lobe (AL) changes with age in the context of alarm pheromone communication in the clonal raider ant (*Ooceraea biroi*) (*17*). Similar to other social insects (*11*, *12*, *16*), older ants responded more rapidly to alarm pheromones, the chemical signals for danger. Using whole-AL calcium imaging (*18*), we then mapped odor representations for five general odorants and two alarm pheromones in young and old ants. Alarm pheromones were represented sparsely at all ages. However, alarm pheromone responses within individual glomeruli changed with age, either increasing or decreasing. Only two glomeruli became sensitized to alarm pheromones with age, while at the same time becoming desensitized to general odorants. Our results suggest that the heightened response to alarm pheromones in older ants occurs via increased sensitivity in these two core glomeruli, illustrating the importance of sensory modulation in social insect division of labor and age-associated behavioral plasticity.

## Results and Discussion

We examined whether clonal raider ants show age-dependent plasticity in alarm behavior using a colony bioassay. Upon stimulation with alarm pheromones, the ants begin to move quickly and disassemble the nest pile where the eggs are located (*17*, *18*). Clonal raider ants can live for up to a year (*19*), so we selected “young ants” (two weeks old) and “old ants” (two months old) to examine effects of age while excluding effects of senescence. Ants were housed in mixed-age colonies and then stimulated with the alarm pheromones 4-methyl-3-heptanone, 4-methyl-3- heptanol, a 9:1 blend of the two based on their ratio found in extracts from ants, or a vehicle control (pentane) (Fig. 1A; Data S1) (*17*). All three alarm pheromone stimuli caused ants to leave the nest in a similar manner, so these samples were pooled for analyzing effects of age (Fig. 1B; see Fig. S1 for these data separately). To determine if age influenced alarm responses, we used nonlinear regression to fit curves to the young and old ant datasets together and separately. For the vehicle control, a single curve for young and old ants was the preferred model (extra sum-of-squares F test), indicating similar baseline behavioral patterns. For the pooled alarm pheromone stimuli, the preferred model incorporated separate curves for young and old ants, with old ants responding faster (Fig. 1C; Table S1). This suggests increased responsiveness to alarm pheromones with age in clonal raider ants, recapitulating prior observations in other ant species and honeybees (*11*, *12*, *16*).

**Figure 1.**
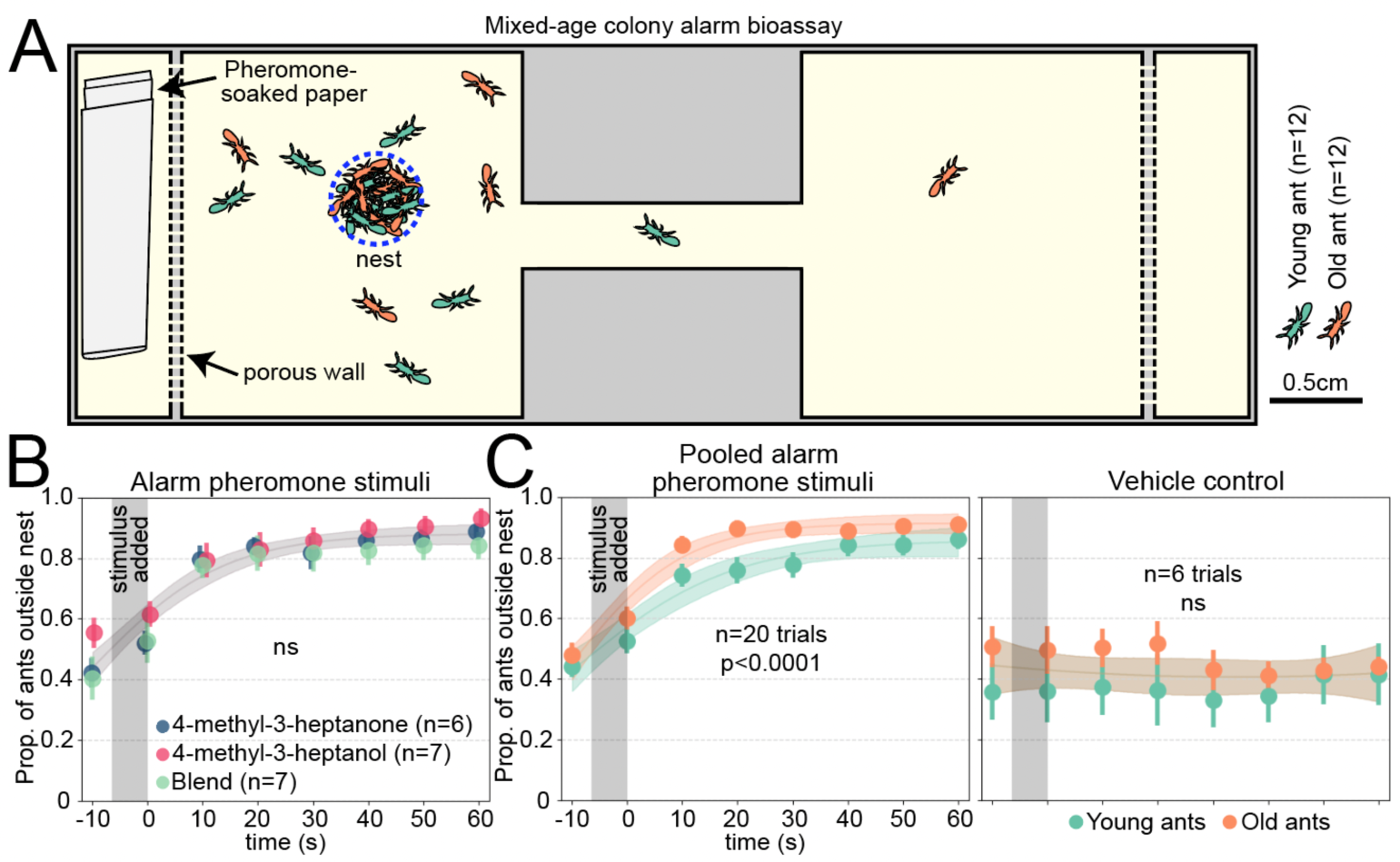
Clonal raider ants exhibit age polyethism in alarm behavior. A) Cartoon of experimental arena. B) Data from young and old ants were pooled, and results from the three alarm pheromone stimuli were compared. Data points are slightly jittered. C) As (B), but showing pooled alarm pheromone trials (left) or vehicle control trials (right), with young and old ants plotted separately and compared statistically. See Fig. S1 for the alarm pheromone stimuli plotted separately. For B&C, we used the extra sum-of-squares F test on same vs. different curves, and vertical grey bars show the time window when the stimulus was added to the arena. See Table S1 for statistical analysis details. Data points show mean±SEM. Ribbons show 95% CI of the curves. ns: p>0.05.

The behavioral differences between young and old ants could result from increased olfactory sensitivity to alarm pheromones, which might be accompanied by broader changes in olfactory function promoting nurse-like vs. forager-like behavioral patterns. Such changes might be detectable by recording odor-evoked calcium responses in the AL, where olfactory sensory neuron (OSN) axons synapse with central brain neurons to form glomeruli (*20*). In a previous study, we used volumetric two-photon microscopy to image whole-AL odor responses in ants expressing GCaMP6s specifically in OSNs (“GCaMP6s ants”) (*18*). Here, we applied the same approach to study putative differences in neural function between young and old ants. Because of the possibility that baseline GCaMP6s signal in the AL might change during aging and therefore affect our ability to detect calcium responses, we quantified AL GCaMP6s fluorescence from dissected brains at both tested ages. Total fluorescence intensity did not differ with age, and calcium responses measured by changes in GCaMP6s fluorescence should therefore be comparable between the two age cohorts (Fig. 2A-B; Data S2).

**Figure 2.**
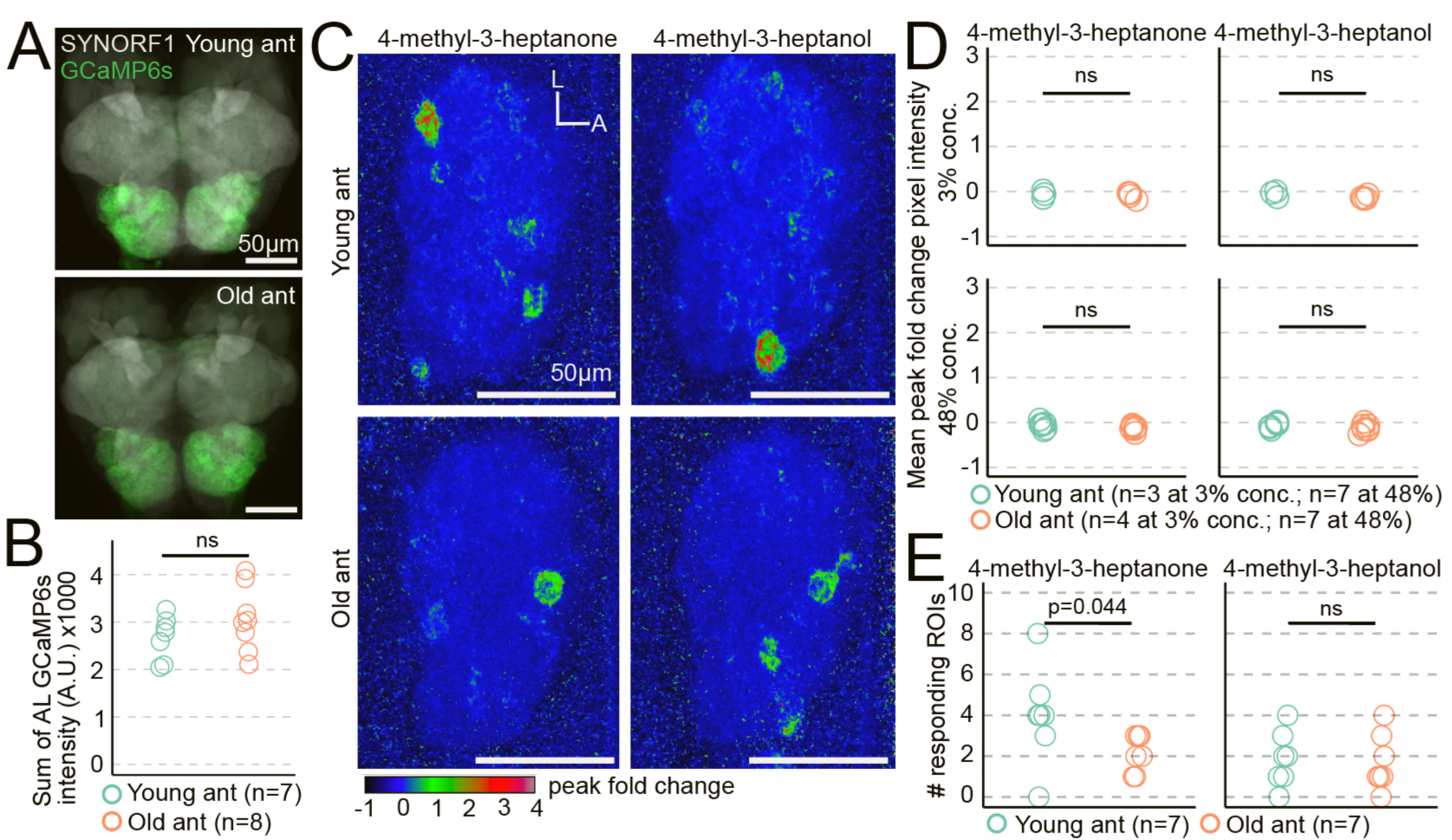
Age-related changes in alarm pheromone representation at the level of the entire antennal lobe. A) GCaMP6s fluorescence in young and old ant brains, stained with anti- SYNORF1 to label neuropil. B) Quantification of AL GCaMP6s fluorescence from young and old ants (p=0.21; Welch’s T-test). C) Exemplar response maps for the two alarm pheromones, showing max Z-projections of the peak fold change across the entire right AL, from a young ant (top) and an old ant (bottom), both at 48% odor concentration v/v. D) Mean pixel intensities across ROIs of the entire AL response maps were compared across age (Welch’s T-tests). E) Comparison of the number of responding ROIs in young ants vs. old ants at 48% odor concentration (Wilcoxon tests). L: lateral; A: anterior. ns: p>0.05.

Next, we imaged AL responses in young and old GCaMP6s ants during stimulation with an odorant panel that evoked robust responses in our prior study at 3% and 48% concentration volume/volume in paraffin oil (alarm pheromones: 4-methyl-3-heptanone and 4-methyl-3- heptanol; general odorants: 3-hexanone, isopropanol, ethylpyrazine, ethanol, and propionic acid) (*18*). Calcium responses to 4-methyl-3-heptanone and 4-methyl-3-heptanol were sparse at both tested ages, and total AL responses did not differ with age for either of the two tested concentrations (Fig. 2C-D; Data S2). We then compared the number of responding ROIs (glomeruli) between the two ages. While there was no significant difference for 4-methyl-3- heptanol, 4-methyl-3-heptanone activated a few more glomeruli in young ants than in old ants (medians of 4 vs. 2 responding ROIs) (Fig. 2E; Data S2). This provides a first indication that alarm pheromone representation changes during aging.

Out of the ∼500 glomeruli in the ant AL, only a small fraction responded to alarm pheromones in young and/or old ants. We therefore asked whether alarm pheromone responses differed quantitatively between age cohorts for these alarm pheromone-sensitive glomeruli. We analyzed two glomeruli that we previously found respond to alarm pheromones in 90 day old ants, “panic glomerulus, broad” (PGb) and “panic glomerulus, alcohol” (PGa) (*18*), as well as four additional glomeruli (G1-G4) in which we reliably observed alarm pheromone responses across individuals (Fig. 3A; Data S2). For G1-G4, we observed a trend toward weaker responses with increasing age, which was statistically significant for responses to 4-methyl-3-heptanone in G2 (Figs. 3B, S2). However, the patterns in PGb and PGa were quite different, with significantly stronger calcium responses in old ants compared to young ants (Figs. 3C-D; PGa does not respond to 4- methyl-3-heptanone, as we reported previously) (*18*). Therefore, while some alarm pheromone-evoked calcium responses decrease during aging, responses in PGb and PGa become elevated instead. The increased sensitivity of PGb and PGa to alarm pheromones with age is consistent with the elevated behavioral response to alarm pheromones in older clonal raider ants (Fig. 1) and increased participation of older individuals in colony defense in other ant species and honeybees (*11*, *12*). A prior study in carpenter ants found only modest effects of age on olfaction as measured by electroantennograms (*7*). This can potentially be explained if sensory responses to the same stimulus can be modulated in opposite directions by different components of the olfactory system. Our findings suggest the intriguing hypothesis that age polyethism in alarm behavior is driven through a re-weighting of olfactory signals within the AL, specifically through increased sensitivity in the PGb and PGa glomeruli.

**Figure 3.**
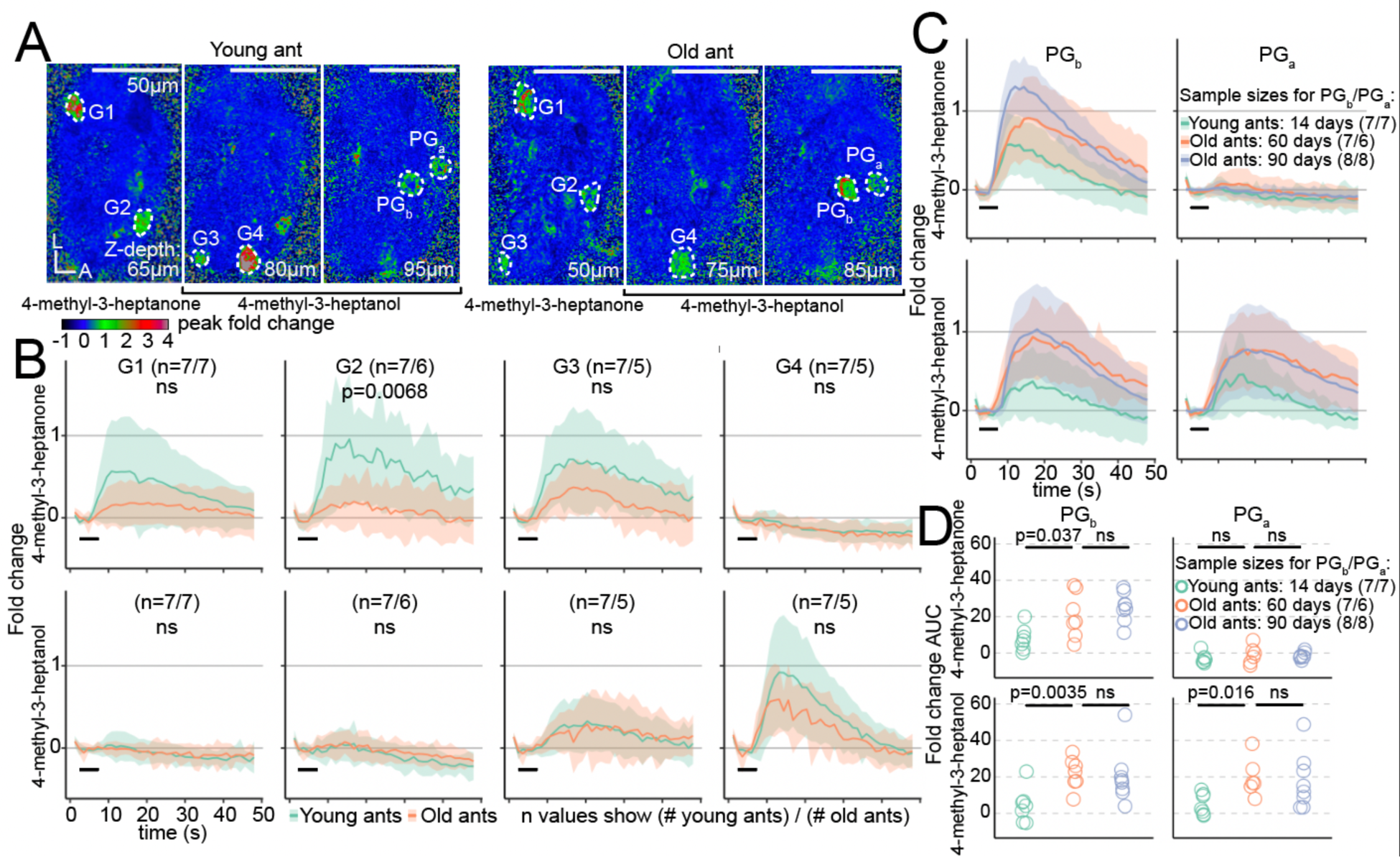
Alarm pheromone representation changes with age at the level of individual glomeruli. A) Calcium responses in selected Z-planes containing the six focal glomeruli in a young ant (left) and an old ant (right). B) Calcium responses to alarm pheromones in focal glomeruli G1-G4. Area under the curve (AUC) values (see Fig. S3) were used to test whether glomerular responses changed with age (Welch’s T-tests). C) Calcium responses to alarm pheromones in PGb and PGa, including data from a previous study on 90 day old ants (*18*). D) AUC from (C). Values were compared across age increments (14 days vs. 60 days, and 60 days vs. 90 days; Welch’s T-tests). Black bars in B&C show the 5s odor stimulation. Time series show mean±SD. Stimuli were presented at 48% concentration v/v. L: lateral; A: anterior. ns: p>0.05.

We then asked whether responses in these glomeruli continue to change later in life by comparing responses to alarm pheromones in PGb and PGa in 60 day old ants with published data from 90 day old ants (*18*). We found no significant differences (Fig. 3C-D; Data S2). Odor codes for alarm pheromones are therefore modulated sometime between 14-60 days of age, and then stably maintained through 90 days of age. This pattern aligns with long-term behavioral tracking experiments in carpenter ants, which found that most workers exhibit either a low-maturity nurse state or a high-maturity forager state, with a fast and permanent switch between states (*5*).

The increased responses to alarm pheromone stimuli in PGb and PGa could represent a specific increase in pheromone sensitivity in these glomeruli or could result from a generalized increase in glomerular excitability. To distinguish between these possibilities, we examined the effects of age on responses to general odorants (Data S2). Strikingly, high concentration stimulus with 3- hexanone and isopropanol frequently generated very broad responses across large portions of the AL in young- but not old ants (Fig. S3A). Mean peak calcium responses across the AL declined dramatically with age for 3-hexanone (>9-fold decline) and isopropanol (>57-fold decline), an effect that was specific to these two odorants when presented at 48% concentration (Fig. S3B). These data suggest that broad-scale changes in olfactory function occur during aging, and that these changes are not limited to alarm pheromone encoding.

We next asked whether any of the general odorants generated excitatory calcium responses in PGb or PGa (Data S2). We found that 3-hexanone activates both glomeruli in both young and old ants, while isopropanol activates both glomeruli in young ants only (Fig. 4A). This result is not surprising given the very broad representation of these two odorants, especially in young ants.

**Figure 4.**
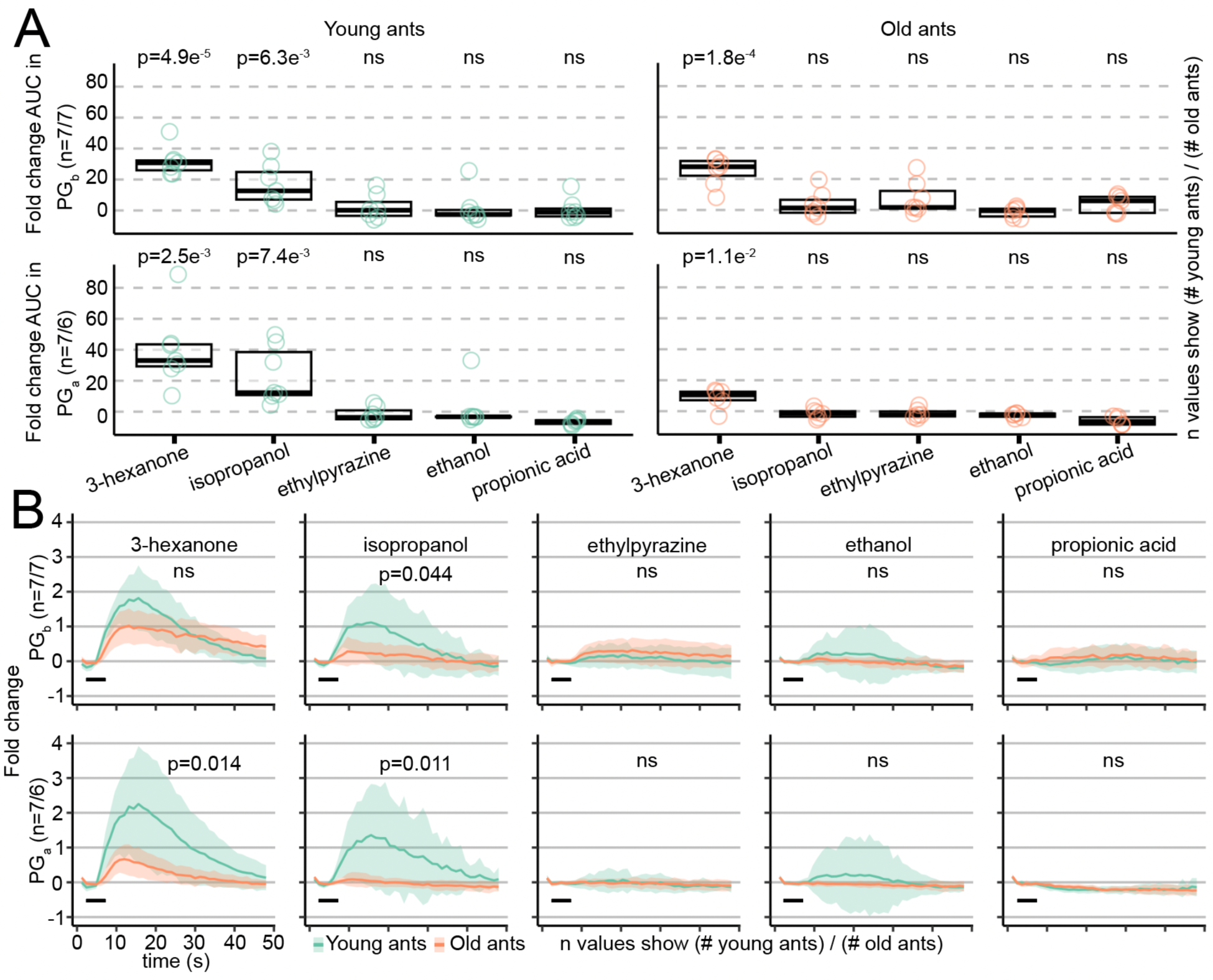
Responses to general odorants in alarm pheromone-sensitive glomeruli are downregulated with age. A) Quantification of odor-evoked responses in PGb (top) and PGa (bottom) in young (left) and old (right) ants. Data show the area under the curve (AUC) of fold change. Boxes contain the first to third quartiles and thick lines show the median. Values were compared against the null expectation of zero excitation (one-sided, one-sample Welch’s T-tests). B) Calcium responses to general odorants in PGb and PGa. AUC values (plotted in Fig. S4A) were used to test whether glomerular responses changed with age (Welch’s T-tests). Black bars show the 5s odor stimulation. Plots show mean±SD. Odors were presented at 48% concentration v/v. ns: p>0.05.

None of the other general odorants activated PGb or PGa at either age (Fig. 4A). Directly comparing calcium responses across age cohorts showed that responses to isopropanol significantly declined with age in both glomeruli, and the same effect was observed for 3- hexanone in PGa (Figs. 4B, S4A). Responses in PGb and PGa to general odorants and pheromones are thus modulated in opposite directions during aging. Interestingly, we also noticed a change in temporal dynamics of calcium responses in one case: responses to 3- hexanone in PGb had a lower peak amplitude but then declined more slowly in old vs. young ants (Figs. 4B, S4B-C). This suggests additional changes in olfactory function occur during aging, beyond simple modulation of response amplitudes. Together, these processes might increase the salience of alarm pheromone stimuli and promote efficient performance of alarm behavior.

There are several possible explanations for the opposite effects of aging on responses to alarm pheromones vs. general odorants in PGb and PGa. Co-expression of odorant receptors and ionotropic receptors in the same OSNs has been documented in flies, mosquitos, and ants (*21–23*), and our results are consistent with a model where PGb and PGa are each associated with two different receptors responsible for detecting general odorants vs. alarm pheromones. The different classes of receptors could be up- or down-regulated in opposite directions during aging (*6*), leading to higher correlation between the presence of alarm pheromone in the local environment and neural activity in the OSNs innervating PGb and PGa. Odor coding is also shaped by interactions between neurons. The basiconic sensilla of the ant antennae can be innervated by dendrites from >100 individual OSNs, and studies in carpenter ants suggest that these neurons are electrically coupled via gap junctions (*24*, *25*). Any age-dependent changes in OSN peripheral connectivity are likely to alter odor responses as measured from AL calcium imaging. In addition, refinement of neuronal architecture within the AL network during adulthood could alter calcium activity in OSN afferents via changing signal input strength or the dynamics of presynaptic modulation (*26–28*). This is in line with evidence for neuroanatomical plasticity over several months of adult life described in other ant species (*27*, *29*), suggesting an extended period of adult neural development.

Odor coding in the ant AL is likely influenced by neuroendocrine and neuromodulator signaling. Juvenile hormone (JH) is a key regulator of insect behavior, including social insect division of labor (*30–32*). Increasing levels of JH with age lead to increased alarm behavior in honeybees without changing antennal responses to pheromones (*11*), while JH signaling reduces the response of AL neurons to aggregation pheromones in locusts (*33*). Various biogenic amines and neuropeptides act on OSN afferents in *Drosophila*, thereby shaping AL odor representation via presynaptic inhibition and facilitation (*34–38*). In ants and other social insects, levels of various neuropeptides in the brain correlate with age and caste, and are implicated in the division of labor (*39–45*). Studies in desert ants detected the neuropeptides allatostatin A and tachykinin within the AL (*46*, *47*), possibly acting on the AL network to regulate odor coding. Additional work is necessary to determine which neuropeptide receptors are expressed on ant OSN afferents, and whether neuropeptide signaling in this region is modulated with age in a manner consistent with our calcium imaging data.

## Conclusions

In this study, we identified age-dependent changes in the neural representation of pheromones and other odorants in the ant AL. While responses to general odorants declined with age in two alarm pheromone-sensitive glomeruli, responses to pheromones were instead upregulated in those same glomeruli, correlating with increased behavioral responses to alarm pheromone stimulation. Our findings highlight the dynamic nature of odor coding in ants across different life stages and suggest an important role for changes in the salience of behaviorally relevant stimuli in generating the division of labor. Olfactory plasticity occurs in many animals, including mammals, and is influenced by factors such as experience, sex, nutrition, and reproductive state (*48–50*). For example, mother mice have increased olfactory bulb responses to behaviorally-relevant natural odors (*51*). Therefore, olfactory modulation is likely an important mechanism that enhances the performance of appropriate behaviors at different life stages across the animal tree of life.

## Acknowledgements

We thank Yohann Chemtob for helpful advice on the behavioral analyses. This work was supported by the National Institute of Neurological Disorders and Stroke of the National Institutes of Health under award number R01NS123899 to D.J.C.K. The content is solely the responsibility of the authors and does not necessarily represent the official views of the National Institutes of Health. This work was also supported by the Howard Hughes Medical Institute, where D.J.C.K. is an investigator. D.D.F. was supported by an Open Philanthropy Fellowship from the Life Sciences Research Foundation. L.E.L. was supported by an NSF Graduate Research Fellowship under award number DGE 194642. This work was supported in part by a grant to The Rockefeller University from the Howard Hughes Medical Institute through the James H. Gilliam Fellowships for Advanced Study program (to L.E.L. and D.J.C.K.). This is Clonal Raider Ant Project paper number 32.

## Author Contributions

T.H. and D.J.C.K. designed the calcium imaging experiments. T.H. performed and analyzed the calcium imaging experiments. L.E.L. designed and performed the behavior experiments and analyzed the data. D.D.F. designed and performed the confocal imaging experiments. T.H. and D.J.C.K. wrote the paper, and all authors read, edited, and approved the manuscript for publication. D.J.C.K. supervised the project.

## Materials & Methods

### Data and Code Availability

- All original code has been deposited to GitHub and is publicly available (https://github.com/Social-Evolution-and-Behavior/Hart_Kronauer2023 (*18*) and https://github.com/Social-Evolution-and-Behavior/Hart_Kronauer2024).
- Any additional information required to reanalyze the data reported in this paper is available from the corresponding authors upon request.

### Ant husbandry and maintenance

Ants were maintained in 5cm Petri dish nests lined with plaster of Paris at 25°C. Colonies were fed frozen fire ant pupae and humidified ∼3 times per week. Ants for behavior experiments were wild type from clonal line B. At the time of behavioral experiments, young ants were 13-19 days post eclosion and old ants were 55-60 days post eclosion. Ants for calcium imaging experiments were from clonal line B and carried the transgene integration [ie-DsRed, ObirOrco-QF2, 15xQUAS-GCaMP6s] (“GCaMP6s ants”) (*18*). GCaMP6s ants were kept in 5cm diameter Petri dish nests and allowed to lay eggs. Eggs were then transferred in batches of ∼50 to a fresh nest and placed with ∼20 wild type adult workers from clonal line A. When the new GCaMP6s ants eclosed to adulthood, the date was recorded and the older wild type workers were removed. All eggs and brood were removed from the experimental GCaMP6s colonies ∼4 days prior to calcium imaging experiments to control the phase of the reproductive cycle at the time of imaging (*52*). When calcium imaging experiments were run, young ants were 15-20 days post eclosion and old ants were 60-61 days post eclosion. To exclude intercastes, only individuals without eyespots were used in calcium imaging experiments (*53*, *54*).

### Behavior

*Chemicals.* 96% 4-methyl-3-heptanone was purchased from Pfaltz and Bauer (Item number M19160). ≥99% 4-methyl-3-heptanol, 98% ethylpyrazine, 99% propionic acid, and 100% pentane were purchased from Sigma-Aldrich (Item numbers M48309, 250384-5G, W292419- SAMPLE-K, and 236705, respectively). 98% 3-hexanone was purchased from Aldrich Chemistry (Item number 103020-10G). 100% ethanol was purchased from Decon Laboratories (Item number 2716), and ≥99.5% isopropanol from Fisher Chemical (Item number A416SK-4). *Mixed-age group alarm bioassay.* The alarm behavior assay was performed as described previously (*17*, *18*), except using clonal line B young and old ants. Individuals were paint tagged to indicate their age. Mixed-age colonies were assembled by placing 12 each of young and old ants in behavioral arenas. Prior to behavioral experiments, ants were allowed to settle for at least 5 days, until they had laid eggs and spent most of their time within a tightly packed nest pile.

Three days after colonies were set, they were supplemented with additional age-controlled ants to replace dead or escaped individuals. From the video recordings, we found that final colony sizes during experiments ranged from 16-24 ants, and the proportion of young ants was 0.49 (SD 0.05). Each colony was tested no more than once per experimental condition.

Each pure compound was freshly diluted 1:200 in 100% pentane each day of experiments. After recording baseline activity for 4 minutes and 30 seconds, 50 µL of each compound was added to a ∼1 cm^2^ piece of filter paper and allowed to evaporate for 30 seconds before folding the paper and placing it into the stimulus chamber. Behavioral responses were recorded for another 5 minutes.

To detect differences in responses to alarm pheromone stimuli between young and old ants, videos were examined to determine when after stimulus presentation ants left the nest. anTraX tracking software was used to identify the spatial coordinates of ants within the colony (*55*). The median location (XY coordinates) of all ants in the first 5 minutes of the video was defined as the center of the nest, and a circle with radius 0.25 cm was defined as the nest. The number of young and old ants outside the nest was counted manually every 10 seconds in the minute prior to and after addition of the stimulus for all videos, with t=0 corresponding to the time when the filter paper was placed in the stimulus chamber of the alarm arena. Ants were considered outside the nest if no part of their body was touching the nest circle. Proportion of ants outside the nest was calculated by dividing the number of young and old ants outside the nest by the total number within the arena.

We focused our statistical analyses on the time 10 seconds prior to addition of the stimulus to 60 seconds after addition of the stimulus. To compare the response of the colonies to the three alarm pheromone stimuli, we fit a logistic growth function with nonlinear regression (least squares) to the total proportion of ants outside the nest in the time of active response to alarm pheromone, and used an extra sum-of-squares F test to ask if a single curve was sufficient to fit the responses to 4-methyl-3-heptanone, 4-methyl-3-heptanol, and the blend, or if the data was better fit with separate curves for each compound. To compare the response of old and young ants within a colony to the alarm stimuli, we fit a logistic growth function with nonlinear regression (least squares) to the old and young ant responses and used an extra sum-of-squares F test to ask if a single curve was sufficient to fit the responses of old and young ants or if the data was better fit by separate curves for old ants and young ants. As the response of old and young ants to the vehicle control could not be fit with a logistic growth function, we used a quadratic (2nd order polynomial) and the same extra sum-of-squares F test approach. Statistical analyses on behavioral data were performed using GraphPad Prism Version 10.0.2 for macOS, GraphPad Software, San Diego, California, USA (https://www.graphpad.com).

### Immunohistochemistry

Antibody staining of ant brains was performed as reported previously (*18*). Briefly, ant brains were dissected in cold phosphate-buffered saline (PBS) and fixed in 4% paraformaldehyde for 2 hours at room temperature. Blocking was performed for at least 2 hours using fresh PBS containing 0.5% Triton X-100 and 5% donkey serum albumin. Samples were incubated with the appropriate dilution of primary antibody (mouse anti-SYNORF1, DSHB #3C11) in fresh blocking solution on an orbital shaker table at room temperature. Following primary incubation, samples were washed and incubated with donkey anti-mouse secondary antibody tagged with Alexa Fluor 647 (Thermo Fisher #A32787) diluted in fresh blocking solution. Stained brains were washed again and mounted in SlowFade Glass antifade mountant (Thermo Fisher #S36917). Young and old ants were dissected and stained in parallel using the same solutions and under identical conditions.

### Confocal Microscopy

Confocal microscopy of antibody-stained brains was conducted using Zen image acquisition software on a Zeiss LSM 900 confocal microscope equipped with 405nm, 488nm, 561nm and 633nm laser lines. Images were obtained using a Zeiss LD LCI Plan-Apochromat 40X / 1.2NA multi-immersion objective lens and Zeiss Immersol G immersion medium (Zeiss # 462959- 9901-000). Z-stacks of whole brains were acquired at 2048x2048 pixel resolution with 1µm Z steps. GCaMP6s intensity in the antennal lobe was quantified using ImageJ/Fiji. Sum Z projections were generated for each brain using the Fiji Z Project function. Rectangular ROIs were then manually drawn around each antennal lobe and the intensity in the GCaMP6s channel quantified using the Fiji Measure function. Young and old brains were imaged using identical laser power and detector gain settings.

### Two photon calcium imaging

All calcium imaging preparations, recordings, stimulus presentations, and image processing were performed as described previously (*18*), and are described here in brief.

#### Specimen preparation

Cold-anesthetized ants were fastened to a custom stage using blue-light curable glue. A layer of parafilm was placed over the ant’s head, and an additional strip of parafilm was used to restrain the antennae ∼1mm in front of an air supply tube. Using a fresh hypodermic needle and sharp forceps, a window was carefully excised through the parafilm and cuticle to reveal the right AL. The ant was perfused with ant saline (127 mM NaCl, 7 mM KCl, 1.5 mM CaCl2, 0.8 mM Na2HPO4, 0.4 mM KH2PO4, 4.8 mM TES, 3.2 mM Trehalose, pH 7.0) (*56*) for the duration of the experiment.

#### Two-photon imaging

The microscope used was a Bruker Investigator with a Coherent Axon 920nm laser. Fluorescent signals were detected through an Olympus 40X 0.9NA water-immersion objective, using a resonant scanning galvanometer, Z-piezo module for high-speed Z-positioning, dual GaAsP detectors, and PrairieView software. In each trial, all GCaMP6s+ glomeruli in the right AL were imaged by scanning a volume of 512x512x33 voxels (XYZ) with 5µm Z steps. Imaging was performed at 2x optical zoom, and the imaging volume spanned 148µm x 148µm x 165µm. Imaging planes were recorded at 27.5 frames per second, with the entire imaging volume scanned every 0.83 seconds. The laser power, detector gain, and position of the imaging volume were regularly re-calibrated so that baseline fluorescence remained visible from all glomeruli. Loss of signal from imaging through tissue was compensated by adjusting the laser power using an exponential function of z-position.

#### Stimulus presentation

Odor stimuli were presented to the ant using a custom-built olfactometer. Filtered air was directed over the antennae at 600mL/min. A 200mL/min ’carrier’ air stream was applied constantly, while a 400mL/min ’stimulus’ air stream was directed through a series of computer-controlled valves. By default, the valves were set to allow stimulus air to combine with carrier air in an odor manifold. During an odor trial, an electronic signal trigger was sent to the valve control system to direct stimulus air through a stimulus vial. The redirection of stimulus air through odor vials had a 3 second delay and lasted for 5 seconds. Stimulus vials were 4mL amber glass vials and contained 300µL of liquid stimulus (either pure paraffin oil as a control, or an odor compound dissolved in paraffin oil). At the beginning of calcium imaging experiments, each ant was stimulated with paraffin oil control stimulus and confirmed to have a constant baseline fluorescence signal before continuing the experiment. Each ant was presented with the seven odor stimuli in a randomized sequence that was repeated three times, for a total of 21 odor stimulus trials. Each ant was presented with odorants only at a single concentration (either 3% or 48% volume/volume dissolved in paraffin oil). Odor stimulus vials were freshly prepared at the beginning of each day of experiments.

#### Image processing

Images were processed using ImageJ/FIJI (*57*). To generate max Z-projections, fluorescence data from each Z-plane was first separated using the ’Deinterleave’ function, and then stabilized using the ’Image Stabilizer’ plugin (*58*). Next, the fluorescence fold change was computed for each Z-plane by subtracting the baseline (the average of frames 1-5, before calcium responses were typically detected) from each frame and then dividing the result by the baseline. The peak fold change was computed by averaging fluorescence fold change from frames 9-14. Then, a 2-pixel minimum filter was applied to each Z-plane to reduce noise using the ’Minimum’ function, and a maximum intensity Z-projection was generated through all Z-planes. Finally, the pixel intensity range was standardized by changing all values >4 to 4 or <-1 to -1 using the “changeValues” function. To calculate the mean peak fold change pixel intensity across the entire AL, an ROI around the boundary of the AL was defined based on a max Z-projection of raw fluorescence.

Responding ROIs were counted by examining all max Z-projections for a given ant, defining a set of ROIs of the approximate size and shape of individual glomeruli that responded in at least two trials, and calculating the mean intensity of the peak fold change for each ROI in all stimulus trials. If the mean of the peak intensity for a given ROI was ≥0.2 when averaged across all trials for a given odorant, that ROI was considered to respond to that odorant. A small number of trials were excluded from this analysis due to motion artifacts that were apparent in the max Z-projection images.

To quantify calcium responses within individual glomeruli, the responding glomeruli were first identified based on their position within the AL, their shape, and their odor response function. Focal glomeruli were chosen because these characteristics were reasonably consistent across individuals, despite the quantitative changes in responsiveness across age categories for some of these glomeruli. Next, the Z-plane containing the center of the responding glomerulus was first identified, and an ROI was created for that glomerulus. Next, a max Z-projection was generated for the chosen Z-plane as well as the Z-planes directly above and below it, and the fluorescence fold change was calculated. The fold change area under the curve (AUC) was then calculated as the sum of the fold change values from frames 6-40. All statistical analyses of calcium imaging data were conducted in R and plotted using ggplot2 (*59*, *60*).

### Quantification and Statistical Analyses

Analyses of behavioral data were performed using GraphPad Prism Version 10.0.2 for macOS. Fluorescence intensity values were statistically compared using R and the packages described and cited above (*59*). Summary data show mean±SD or another format as indicated in the relevant figure legends.

## Supplemental Information Titles and Legends

**Data S1. Behavioral data.** CSV file containing numbers of young and old ants inside the behavior arena, and proportions of ants of each age outside the nest throughout the assay. The “filename” column indicates the trial, with “rd#_arena#” specifying colony ID.

**Data S2. Calcium imaging quantifications.** Excel file containing quantifications of brain imaging data. Sheet 1 “Total AL GCaMP fluorescence” is related to Fig. 2B. Sheet 2 “Whole-AL peak fold change” is related to Figs. 2D and S3B. Sheet 3 “Number responding ROIs” is related to Fig. 2E. Sheet 4 “Time series in focal glomeruli” is related to Figs. 3, 4, S2, and S4.

## Supplemental Figures and Tables

**Figure S1.**
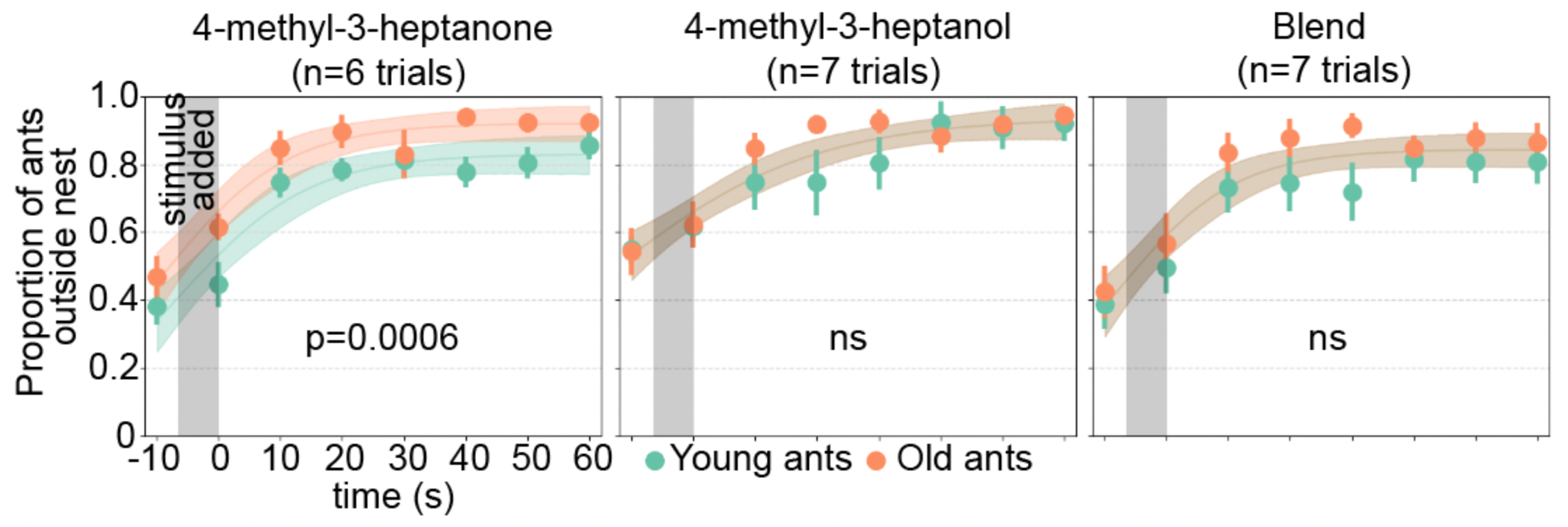
Colony alarm bioassay data for individual alarm pheromone stimuli. Data points show mean±SEM. Raw data were fitted with logistic regressions, with ribbons showing 95% CI. P values relate to whether the single curve model was rejected in favor of separate curves for young and old ants (extra sum-of-squares F test). For 4-methyl-3-heptanone, a model with separate curves was preferred. For 4-methyl-3-heptanol and the blend, similar trends were observed, but single curve models were preferred. See Table S1 for details on the statistical analyses. Grey bars show the time window when the stimulus was added to the arena. ns: p>0.05.

**Figure S2.**
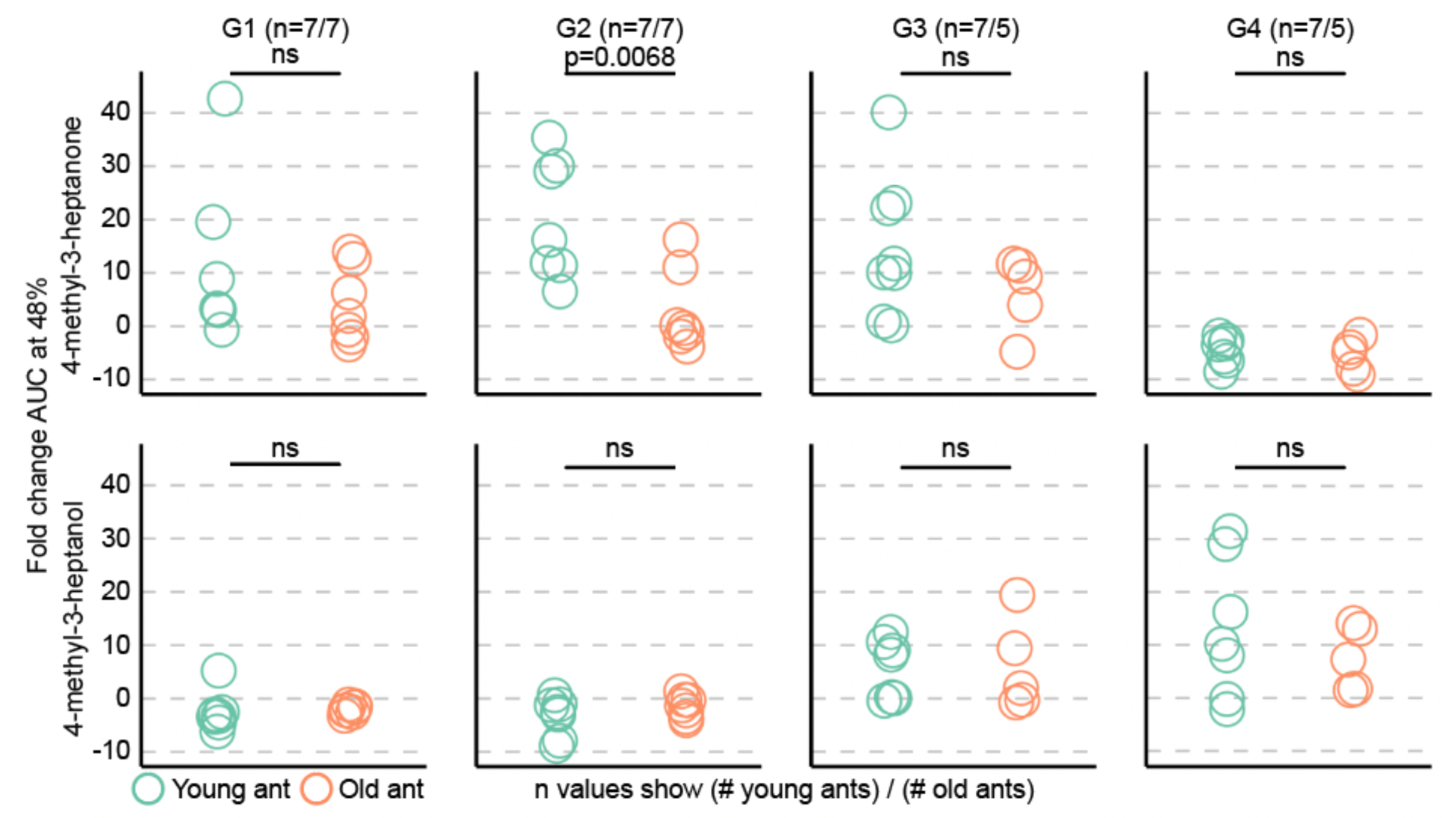
Additional quantification of alarm pheromone responses in focal glomeruli G1- G4. Area under the curve values calculated from the calcium response time series data shown in Fig. 3B. Statistical test results are shown here again (Welch’s T-tests). ns: p>0.05.

**Figure S3.**
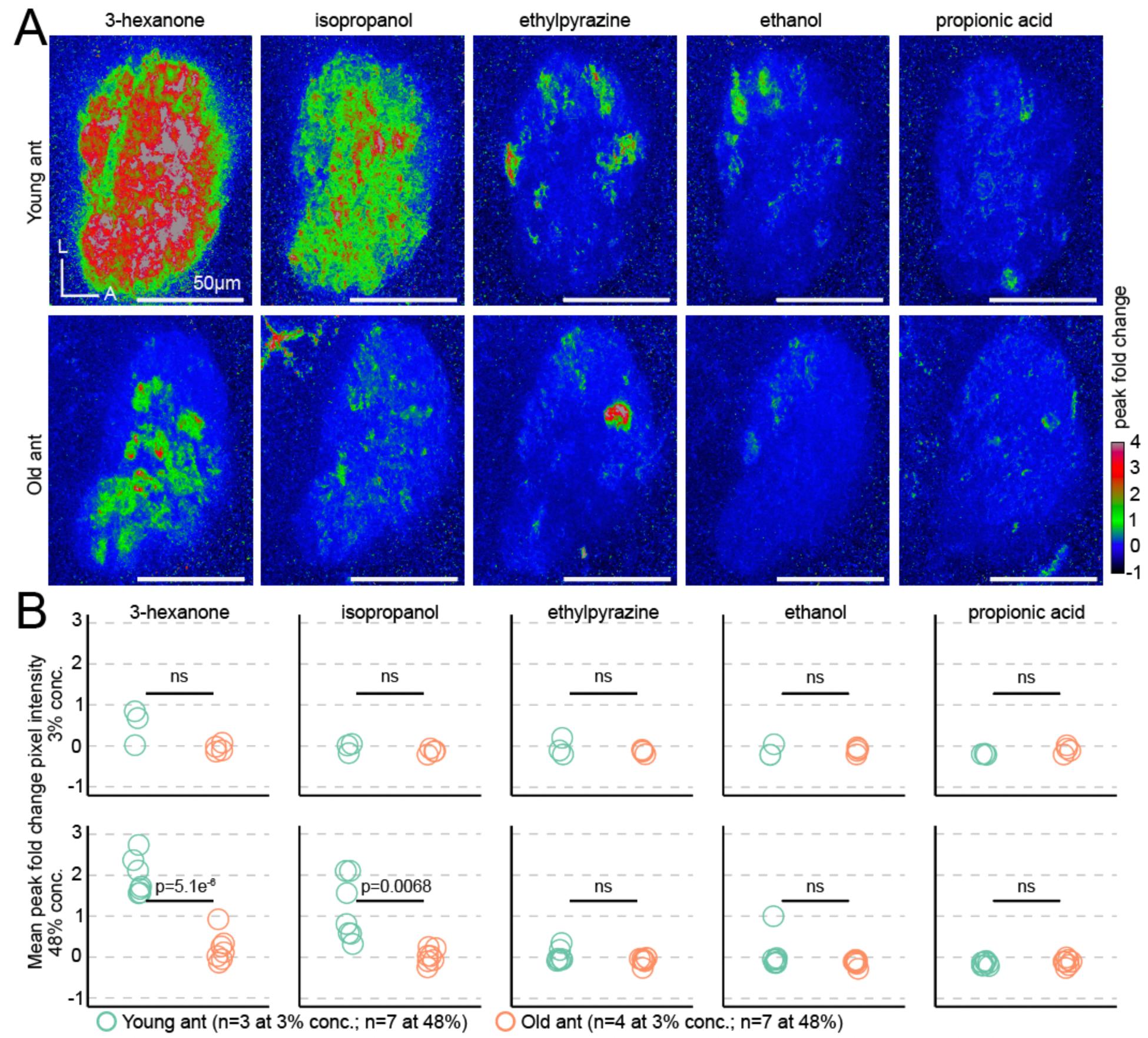
General odorant responses are modulated with age. A) Exemplar response maps for the five general odorants, showing max Z-projections of the peak fold change across the entire right AL, from a young ant (top) and an old ant (bottom), both at 48% odor concentration v/v. B) Mean pixel intensities across ROIs of the entire AL response maps, at 3% (top) and 48% odor concentration (bottom). Responses were compared across age (Welch’s T-tests). L: lateral; A: anterior. ns: p>0.05.

**Figure S4.**
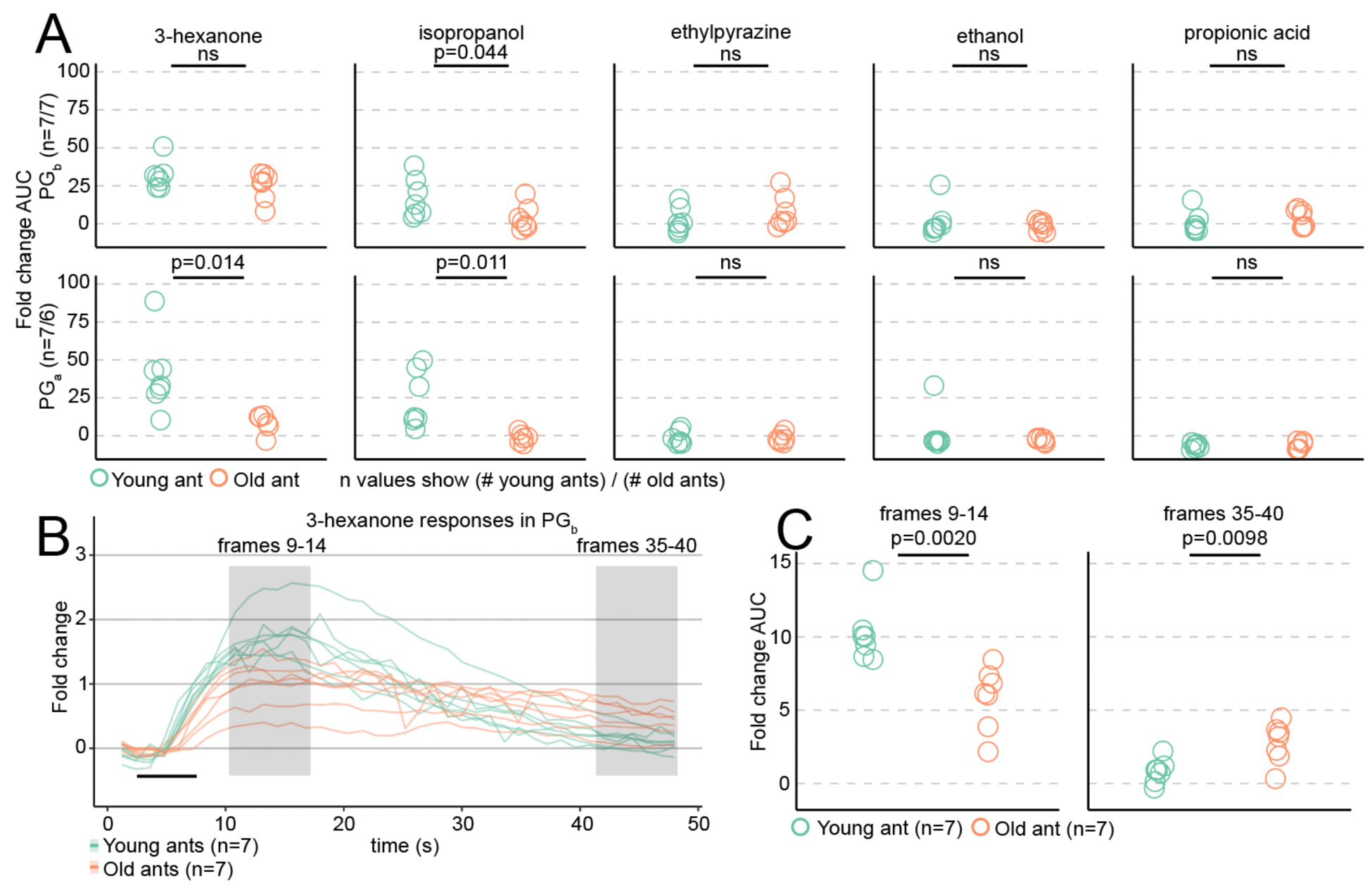
Additional quantification of general odorant responses in PGb and PGa. A) Area under the curve (AUC) of the time series data shown in Fig. 4B; effects of age were tested (Welch’s T-tests). B) Calcium response time series in PGb from individual ants stimulated with 3- hexanone at 48% concentration v/v, to show the effect of age on temporal dynamics of the response. Each line shows the mean of three trials from the same ant. Two time windows of interest are highlighted with grey rectangles. C) Mean AUC values were calculated from the two time windows highlighted in (B). For frames 9-14, containing the peak of the response, AUC values were significantly greater in 14 day old (“young”) ants compared to 60 day old (“old”) ants (left, Welch’s T-test). For the final frames of the recordings (frames 35-40), the opposite relationship was found (right, Welch’s T-test). ns: p>0.05.

**Table S1.**
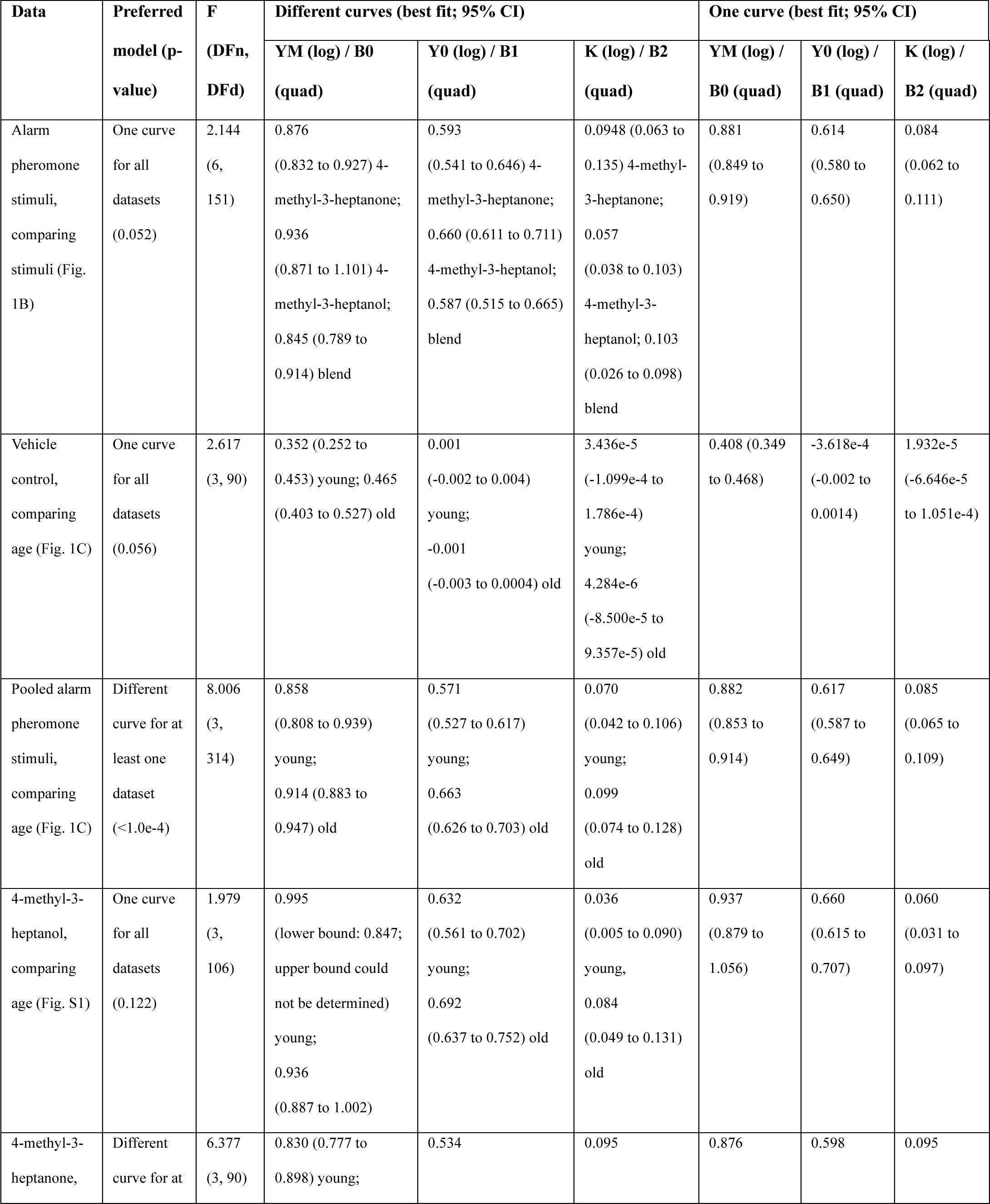
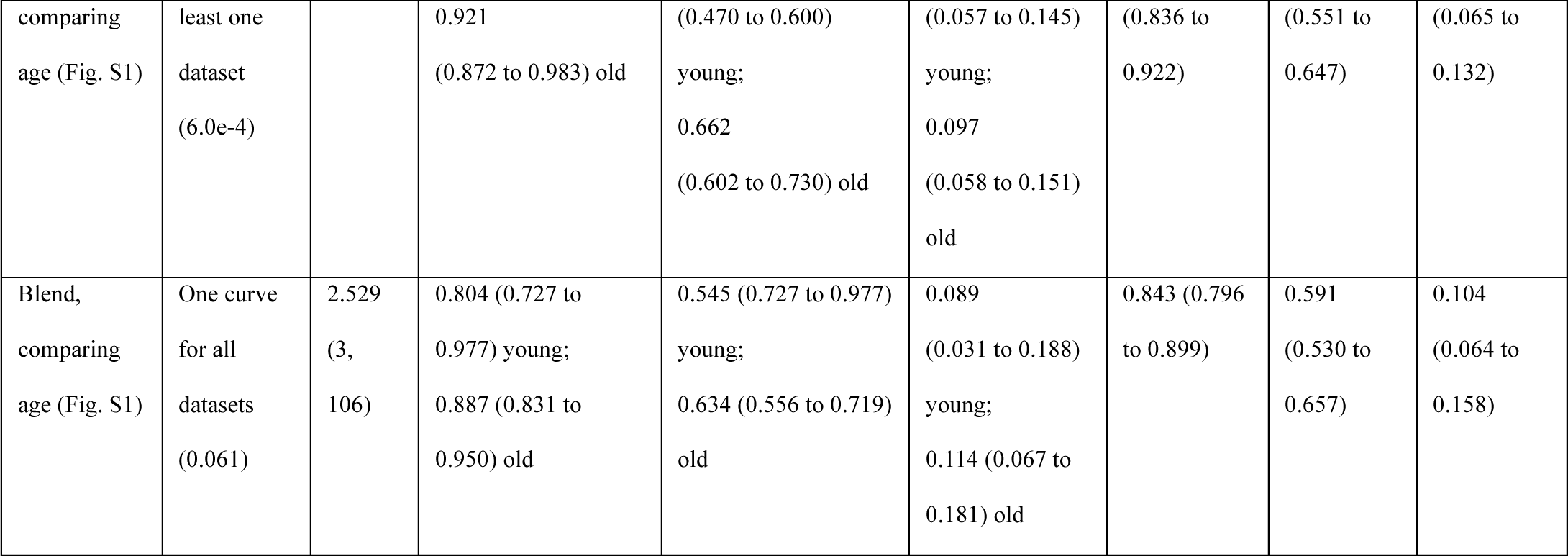
Statistical analyses of behavior experiments. The table includes comparisons of nonlinear regression models when fitting separate curves to each dataset or a single curve to all datasets. The preferred model was determined using the extra sum-of-squares F test. Logistic growth functions (Y=YM*Y0/((YM-Y0)*exp(-k*x)+Y0) were fit to all datasets except for the vehicle control, where a quadratic function was fit (Y=B0+B1*X+B2*X^2^).

## Notes

### Competing Interest Statement

The authors have declared no competing interest.

